# A commercial autogenous injection vaccine protects ballan wrasse (*Labrus bergylta*, Ascanius) against *Aeromonas salmonicida vapA* type V

**DOI:** 10.1101/2020.07.02.183616

**Authors:** J. Gustavo Ramirez-Paredes, D. Verner-Jeffreys, A. Papadopoulou, S. J Monaghan, L. Smith, D. Haydon, T. S. Wallis, A. Davie, A. Adams, H. Migaud

## Abstract

Atypical *Aeromonas salmonicida* (a*As*) and *Vibrionaceae* related species are bacteria routinely recovered from diseased ballan wrasse used as cleaner fish in Atlantic salmon farming. Autogenous multivalent vaccines formulated from these microorganisms are widely used by the industry to protect farmed wrasse despite limited experimental proof that they are primary pathogens. In this study, the components of a commercial multivalent injection wrasse vaccine were tested for infectivity, pathogenicity and virulence via intra peritoneal injection at pre-deployment size (25-50g) and the efficacy of the vaccine for protection against a*As* assessed. Injection with 3.5×10^9^, 8×10^9^ 1.8×10^9^ and 5×10^9^ cfu/fish of *Vibrio splendidus*, *V. ichthyoenteri*, *Aliivibrio logeii* and *A. salmonicida*, respectively, did not cause significant mortalities, lesions or clinical signs after a period of 14 days. IP injection with both a*As* and *Photobacterium indicum* successfully reproduced the clinical signs and internal lesions observed during natural outbreaks of the disease. Differences in virulence (LD_50_ at day 8-post infection of 3.6×10^6^ cfu/fish and 1.6×10^7^ cfu/fish) were observed for two a*As vapA* type V isolates. In addition, the LD_50_ for *Photobacterium indicum* was 2.2×10^7^ cfu/fish. The autogenous vaccine was highly protective against the two a*As vapA* type V isolates after 700-degree days of immunisation. The RPS_FINAL_ values for the first isolate were 95 and 91% at 1×10^6^ cfu/fish and 1×10^7^ cfu/fish, respectively, and 79% at 1×10^7^ cfu/fish for the second isolate tested. In addition, significantly higher anti a*As* seral antibodies (IgM), were detected by ELISA in vaccinated fish in contrast with control (mock vaccinated) fish. These results suggest wrasse can be effectively immunised and protected against a*As* infection by injection with oil adjuvanted vaccines prepared with inactivated homologous isolates. Further work should assess the efficacy of vaccination against other isolates that have proven to be pathogenic such as a*As* type VI and *Photobacterium indicum* and explore the feasibility of immersion vaccination. In addition, a full characterisation of a*As* isolates within the same *vapA* types should be performed as differences in virulence between *vapA* type V isolates were observed and partial genome analysis indicated small but potentially important genomic differences in these isolates.

## INTRODUCTION

Bacterial pathogens are considered the major cause of infectious diseases and mortalities in farmed ballan wrasse produced for sea lice control in the salmon farming industry [1–3]. In Scotland, atypical *Aeromonas salmonicida (*a*<u>As</u>) vapA* type V and VI*, Vibrio splendidus*, *V. ichthyoenteri*, *Aliivibrio salmonicida*, *A. logeii*, and *Photobacterium indicum* are the bacterial pathogens most frequently isolated from ballan wrasse during outbreaks of disease in both hatcheries and post deployment in salmon sea sites [4]. Similar reports are available for the species in Norway [3].

Immunisation of farmed salmonids (Atlantic salmon and rainbow trout) against typical *As* using fully licenced oil-adjuvanted injectable vaccines has historically proven successful and is a standard practice [5]. However, immunisation of non–salmonid species against typical and atypical *As* has been rather challenging [6–8]. For instance, an experimental vaccine containing atypical strains protected Arctic charr (*Salvelinus alpinus*, L.) but not in European grayling (*Thymallus thymallus*, L.) [9]. Furthermore, commercial furunculosis vaccines for salmonids have induced protection in Atlantic halibut (*Hippoglossus hippoglossus*, L.) but not in Atlantic cod (*Gadus morhua*, L.) or turbot (*Scophthalmus maximus*, L.) [6–8].

Currently no licenced or registered vaccines are commercially available in the UK for the prevention and control of infectious diseases in ballan wrasse. Therefore, prophylactic treatments in Scotland are mainly based on the use of autogenous vaccines, which are formulated with antigens derived from pathogens recovered during episodes of elevated mortality [2].

Autogenous or “herd specific” vaccines are farm specific immunological veterinary medicinal products that have the potential to be rapidly developed and deployed when no off-the shelf fully licensed vaccines exist or these have proven infective. In principle, autogenous vaccines must be inactivated (killed), manufactured in licenced facilities, used only under veterinary prescription and on the sites where the pathogens were isolated [10–13].

In Scotland, autogenous vaccines for ballan wrasse were first developed from isolates collected during disease outbreaks between 2013 and 2014 (Ridgeway Biologicals Ltd.) and used in hatcheries and wild caught wrasse. The vaccine formulation later evolved and new isolates were introduced following a health screening surveys [4]. However without established challenged models for the Scottish bacteria and wrasse populations, the actual virulence of the isolates and the efficacy/potency of the vaccine components remained unknown.

Overall, i.p. injection challenges with atypical strains of *As* have been successful in several species with a wide range of doses used [14]. For instance, juvenile spotted wolfish (*Anarhichas minor*, L) succumbed to disease when i.p. injected with a*As* at 10^3^ and 10^4^ cfu / mL [8, 15], while high morbidities in turbot were reported [16], but only in fish exposed with the same method to 10^8^ and 10^10^ cfu / mL. Experimentally infected ballan wrasse and lumpsucker also experienced high morbidities (> 70%) when challenged with Norwegian a*As* isolates at doses of 2 ×10^3^ cfu / mL (bath) and 2 × 10^6^ cfu /mL (i.p. injection), and 10^8^ cfu /mL (i.p. injection), respectively [17, 18]. As for the *Vibrionaceae* pathogens in cleaner fish, in a previous study in Norway, only *Vibrio anguillarum* originally isolated from Atlantic salmon caused high mortalities (up to 60%) in ballan wrasse under experimental conditions, while Norwegian ballan wrasse isolates of the same bacterial species caused < 20% mortalities when challenged via bath, cohabitation and i.p. injection [17].

Given that, a*As* and *Vibrionaceae* isolates are highly heterogenic and variable, and virulence is often strain and host dependant [1, 19], the establishment of similar experiments in other geographical areas such as Scotland is of high relevance for the local industry.

In the present study, *in vivo* challenge models were developed via intraperitoneal (i.p.) injection in Scottish ballan wrasse (25-50 g) to investigate the infectivity, pathogenicity and virulence of isolates routinely recovered from diseased wrasse and used as antigens in commercial autogenous vaccines. These isolates included a*As vapA* types V and VI, *Vibrio splendidus*, *V*. *ichthyoenteri*, *Aliivibrio salmonicida*, *A*. l*ogeii*, and *Photobacterium indicum*. Furthermore, the efficacy of the a*As vapA* type V components of the vaccine was assessed by measuring survival rates after experimentally infecting vaccinated and control fish by i.p. injection with homologous isolates at medium, high and very high doses and its potency expressed in terms of RPS. Specific antibody (IgM) kinetics were assessed as a relevant correlate of protection.

## MATERIALS AND METHODS

### Bacterial identification and genotyping

The bacterial isolates used were recovered from diseased fish at commercial hatcheries and characterised on the basis of phenotypic and genotypic characteristics as part of a previous study [4]. In brief, bacterial DNA was extracted using Genesig® Easy DNA/RNA Extraction Kit (Primerdesign Ltd, Southampton UK) according to the manufacturer’s instructions. Species confirmation was performed on the samples by targeting the V3-V4 hypervariable region of the *16S rRNA* gene [20] and the subunit B protein of DNA gyrase (topoisomerase type II) – *gyrB* gene [21]. The *Aeromonas salmonicida* isolates were then genotyped by sequencing the A-layer membrane as described previously [22].

For the experimental infections, the a*As* isolates were grown on tryptone soya agar (TSA, Oxoid, UK) or blood agar (BA; TSA + 5% sheep blood Thermo Fisher) while the *Vibrionaceae* isolates were on sea water agar (SWA, Oxoid, UK) and incubated at 22 °C for 48 and 24 h, respectively. For growth in liquid media, a*As* isolates were inoculated onto trypticase soy broth (TSB, Oxoid, UK) and *Vibrionaceae* isolates onto TSB + 2% NaCl (Oxoid, UK,) and incubated at 22 °C for 18-24 h, with continuous shaking at 180 rpm. For harvesting, all bacteria were centrifuged at 4 °C for 10 min at 2,000 x g, bacterial pellets were then washed with sterile 1x phosphate-buffered saline (PBS) and resuspended in sterile PBS to the required concentration (cfu/mL) for the experiments.

With the exception of isolate TW164/15 (a*As vapA* type VI) that was recovered from moribund lumpsucker (*Cyclopterus lumpus*) the rest of the isolates were recovered from ballan wrasse. A summary of the isolates used in this study is presented in Table 1.

**Table 1.**
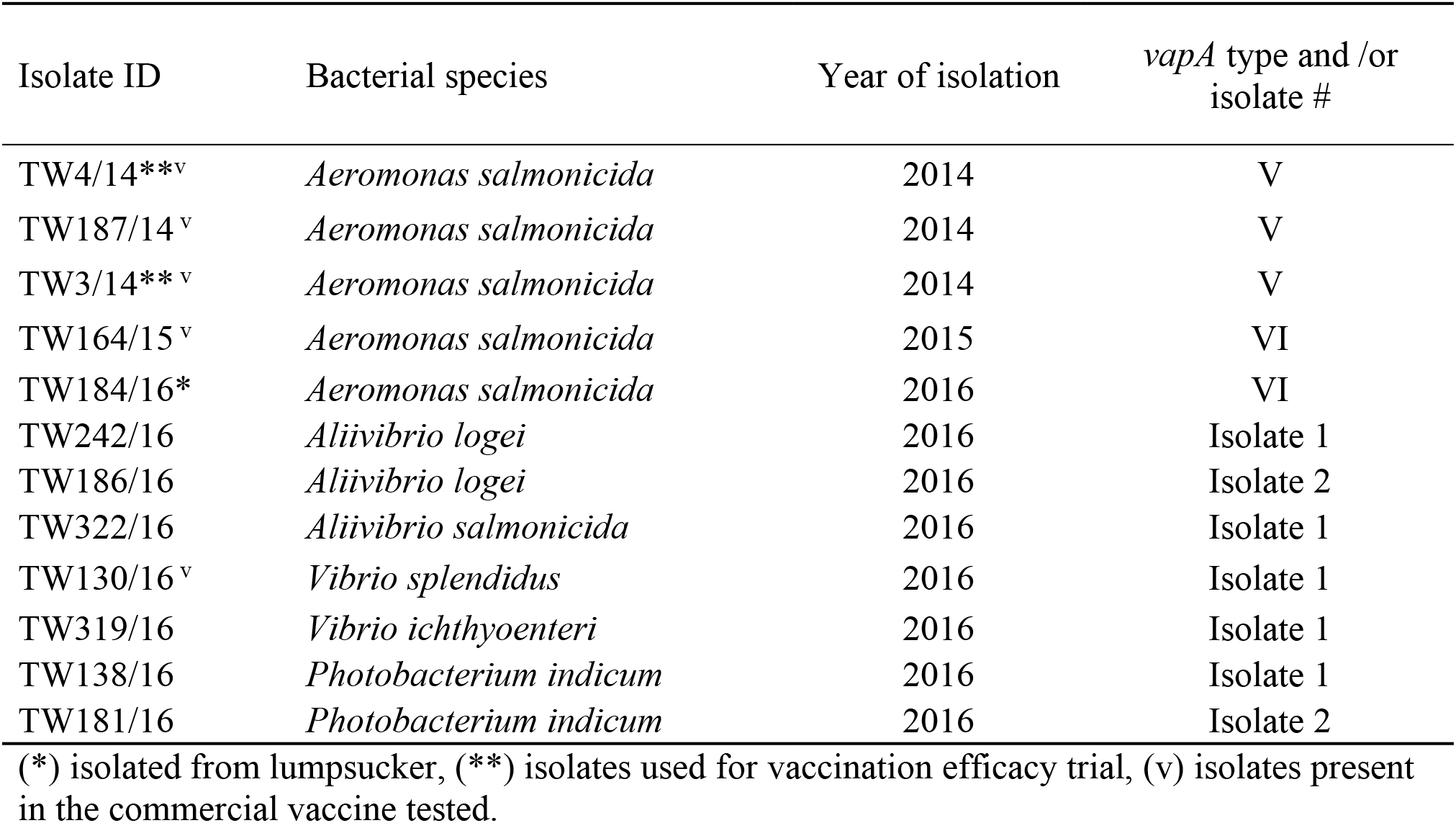
Bacterial isolates used in this study for pathogenicity, virulence and vaccine assessment.

### Experimental fish

A population of naïve *i.e.* unvaccinated and non-diseased ballan wrasse (30 ± 5 g) was provided by a commercial cleaner fish hatchery on the west coast of Scotland. Prior to the study, the health status of the fish was checked by screening a subset of the population with standard histological, bacteriological and molecular methods to confirm the absence of a*As* [4, 22], amoebic gill disease [23] and *Vibrionaceae* related bacteria [20, 21]. After confirmation they were free of these pathogens, fish were transferred to the Centre for Environment, Fisheries and Aquaculture Science (Cefas) Weymouth Laboratory in February 2017.

Fish were acclimated and quarantined for 3 weeks after arrival in 6 aerated aquaria (approx. 900 L, tanks enriched with artificial plastic kelp and sections of plastic pipes to provide hides to the fish) at 12.0 ± 0.5 °C with a 20:4 h light:dark photoperiod, water flow of 4.0 L / min and dissolved oxygen (DO) at 8 ± 0.5 mg / L. During this period, fish were further screened for bacteriology (swabs from head kidney plated onto SWA), histopathology (fixed in 10% neutral buffered formalin) and molecular methods as described before. In addition, virology diagnostic tests were performed to discard the presence of notifiable viral diseases as per the protocols in the OIE manual of diagnostic tests for aquatic animals [24].

### Vaccine

A commercial injectable (oil-based) multivalent autogenous vaccine, containing antigens from isolates TW3/14, TW4/14, TW187/14, TW164/15 and TW130/16 (Table 1) was provided by Ridgeway Biologicals Ltd. The vaccine was shipped to Cefas and stored at 4 ± 1 °C prior to use.

### Confirmation of infectivity of components of multivalent autogenous vaccine

The virulence of bacterial isolates, representative of strains commonly used as components of the multivalent autogenous vaccines used in the industry, was assessed in a series of infection experiments performed in 30 L tanks enriched as for acclimation tanks (Table 2).

**Table 2.** Bacterial isolates, number of fish and doses used in the different infection experiments to assess infectivity, pathogenicity and virulence.

| Infection Experiment | Number of fish i.p. injected | Days observation post challenge | Bacterial species and isolate | OD<sub>600</sub> / dilution factor | Dose (cfu/ fish) | No. dead/moribund by termination |
| --- | --- | --- | --- | --- | --- | --- |
| Experiment 1 | 6 | 6 | *aAs* type V – TW4/14 | 1.46 | 9.5 x 10<sup>7</sup> | 6 (100%) |
|  | 6 | 6 | *aAs* type V – TW187/14 | 1.52 | 8.5 x 10<sup>7</sup> | 6 (100%) |
|  | 6 | 6 | *aAs* type V – TW3/14 | 1.49 | 1.0 x 10<sup>8</sup> | 6 (100%) |
|  | 6 | 6 | *Aliivibrio salmonicida* TW322/16 | 1.51 | 5.0 x 10<sup>4</sup> | 0 |
|  | 6 | 6 | *Vibrio splendidus* TW130/16 | 1.45 | 2.0 x 10<sup>5</sup> | 0 |
|  | 6 | 6 | *Vibrio ichthyenteri* TW319/16 | 1.48 | 1.0 x 10<sup>9</sup> | 0 |
| Experiment 2 | 15 | 7 | *Aliivibrio salmonicida* TW322/16 | >2.5 | 5.0 x 10<sup>9</sup> | 2 (13%) |
|  | 15 | 7 | *Vibrio splendidus* TW130/16 | >2.5 | 3.5 x 10<sup>9</sup> | 0 |
|  | 15 | 7 | *Vibrio ichthyenteri* TW319/16 | >2.5 | 8.0 x 10<sup>9</sup> | 0 |
|  | 15 | 7 | Control - 1x PBS | - | - | 0 |
|  | 15 | 8 | *aAs* type VI – TW164/15 | 2 | 3.0 x 10<sup>9</sup> | 15 (100%) |
| Experiment 3 | 12 | 15 | *aAs* type VI – TW164/15 | 2 | 3.0 x 10<sup>9</sup> | 12 (100%) |
|  | 12 | 15 | *aAs* type VI – TW164/15 | 10<sup>-2</sup> | 3.0 x 10<sup>7</sup> | 12 (100%) |
|  | 12 | 15 | *aAs* type VI – TW184/16 | 2 | 1.6 x 10<sup>9</sup> | 12 (100%) |
|  | 12 | 15 | *aAs* type VI – TW184/16 | 10<sup>-2</sup> | 1.6 x 10<sup>7</sup> | 12 (100%) |
|  | 12 | 12 | *Photobacterium indicum* TW138/16 | 1.75 | 3.2 x 10<sup>9</sup> | 12 (100%) |
|  | 12 | 12 | *Photobacterium indicum* TW138/16 | 10<sup>-2</sup> | 3.2 x 10<sup>7</sup> | 6 (50%) |
|  | 12 | 12 | *Photobacterium indicum* TW181/16 | 1.89 | 9.0 x 10<sup>9</sup> | 12 (100%) |
|  | 12 | 12 | *Photobacterium indicum* TW181/16 | 10<sup>-2</sup> | 9.0 x 10<sup>7</sup> | 8 (66%) |
|  | 12 | 25 | *Aliivibrio logei* TW242/16 | 2 | 1.5 x 10<sup>9</sup> | 0 |
|  | 12 | 25 | *Aliivibrio logei* TW242/16 | 10<sup>-2</sup> | 1.5 x 10<sup>7</sup> | 0 |
|  | 12 | 25 | *Aliivibrio logei* TW186/16 | 2 | 1.8 x 10<sup>9</sup> | 0 |
|  | 12 | 25 | *Aliivibrio logei* TW186/16 | 10<sup>-2</sup> | 1.8 x 10<sup>7</sup> | 0 |
| Experiment 4 | 15 | 16 | *aAs* type V – TW4/14 | 1.9 | 1.0 x 10<sup>8</sup> | 15 (100%) |
|  | 15 | 16 | *aAs* type V – TW4/14 | 10<sup>-1</sup> | 1.0 x 10<sup>7</sup> | 13 (87%) |
|  | 15 | 16 | *aAs* type V – TW4/14 | 10<sup>-2</sup> | 1.0 x 10<sup>6</sup> | 8 (53%) |
|  | 15 | 16 | *aAs* type V – TW4/14 | 10<sup>-4</sup> | 1.0 x 10<sup>4</sup> | 0 |
|  | 15 | 16 | Control - 1x PBS | - | - | 0 |
|  | 15 | 19 | *aAs* type VI – TW164/15 | 2 | 2.5 x 10<sup>8</sup> | 1 (7%) |
|  | 15 | 19 | *aAs* type VI – TW164/15 | 10<sup>-1</sup> | 2.5 x 10<sup>7</sup> | 0 |
|  | 15 | 19 | *aAs* type VI – TW164/15 | 10<sup>-2</sup> | 2.5x 10<sup>6</sup> | 0 |
|  | 15 | 19 | *aAs* type VI – TW164/15 | 10<sup>-4</sup> | 2.5 x 10<sup>4</sup> | 0 |
|  | 15 | 19 | Control - 1x PBS | - | n/a | 0 |
| Experiment 5 | 15 | 19 | *aAs* type V – TW4/14 | 1.95 dil. 10<sup>-1</sup> | 1.0 x 10<sup>7</sup> | 15 (100%) |
|  | 15 | 19 | *aAs* type V – TW4/14 | 10<sup>-2</sup> | 1.0 x 10<sup>6</sup> | 8 (53%) |
|  | 15 | 19 | *aAs* type V – TW4/14 | 10<sup>-3</sup> | 1.0 x 10<sup>5</sup> | 7 (47%) |
|  | 15 | 19 | *aAs* type V – TW4/14 | 10<sup>-4</sup> | 1.0 x 10<sup>4</sup> | 1 (7%) |
|  | 15 | 19 | *aAs* type V – TW4/14 | 10<sup>-5</sup> | 1.0 x 10<sup>3</sup> | 0 |

For the first infection experiment, limited numbers of 30 ± 5 g fish (n= 6) were injected with an OD_600_ ~1.57 bacterial suspension of different isolates representing 4 different bacterial species (a*As* including two type V and one type VI isolates, *Vibrio splendidus*, *Aliivibrio salmonicida* and *Vibrio ichthyoenteri*) (Table 2).

In the second infection experiment, the pathogens that did not cause morbidities or signs of disease during the first infection experiment were i.p. injected in naïve ballan wrasse at a higher dose. To confirm that these isolates were not pathogenic via this exposure route, the number of fish tested was also increased to 15 per isolate, and the length of the experiment was prolonged to 16 days. In addition, an a*As vapA* type VI (isolate TW164/15) recovered from lumpsucker was also included (Table 2).

In the third infection experiment, fish (n= 12) were i.p. injected with medium (10^7^ cfu/fish) and high (10^9^ cfu/fish) doses of 2x isolates of *Aliivibrio logei* and *Photobacterium indicum* as well 2x isolates of a*As vapA* type VI and observed for at least 25 days (Table 2). The isolates used were prepared directly from cryopreserved stocks and had not previously passaged in fish.

Moribund fish and mortalities from all experiments were removed from the tanks, their external and internal condition assessed. Head kidney swabs were taken onto solid media for bacteriological assessment. Isolates not recovered, despite being i.p injected into the fish at high doses, were regarded as non-infectious. The bacteria recovered were subcultured to purity, their identities confirmed and cryopreserved at – 80 °C until further use.

Additional infection experiments 4 and 5 were also undertaken. These were to better determine both the relevant virulence of the different a*As* isolates *vapA* type V and identify doses that would ideally result in high, but not excessive (50-75% mortality), suitable for use in vaccine efficacy testing. In infection experiment 4, 4x different doses of each pathogen were tested (n= 15 fish per dose) with a control treatment (PBS) included. Initial results generated by infection experiment 4 were confirmed in a second set of pre-tests with a longer observation period post injection (4 weeks) without PBS controls (Table 2). The isolates used were passaged (recovered from moribund fish) from infection experiments 1 and 2 (Table 2).

For the isolates where the use of lower and higher doses caused a mortality response below and above 50% respectively, the median lethal dose (LD_50_) was calculated according to [27] to define and compare their virulence at the time point of occurrence. Results obtained from both experiments 4 and 5 were used to select isolates for vaccine testing and determine the doses for the main challenge infection in the vaccine efficacy trial (Table 2). In addition, differences within the a*As vapA* type V isolates TW4/14, TW187/14 and TW3/14 were investigated with macrorestriction analysis using pulsed field gel electrophoresis (PFGE) as described previously [25] with the following modifications. Bacteria were grown on TSA at 15 °C for 72 – 96 h, *SpeI* restriction enzyme (5U per 150 μL, New England Biolabs) was used [26] and the electrophoresis conditions comprised switch times of 2 – 6 s at 15 °C and 200 V for 37 h.

For all the infection experiments, fish were transferred from a stock tank, anaesthetised with MS-222 (40 ppm; Tricaine methane sulphonate, Sigma) and i.p. injected with 100 μL of the relevant bacterial suspension. Where included, control fish were injected with 100 μL of sterile PBS. Fish were then allocated to respective 30 L aquaria each with water flow of 0.6 – 1.0 L / min, all other parameters remained the same as described above. Fish were observed at least twice a day for signs of disease for 7-14 days. The pathogens that caused mortalities, were recovered from the diseased fish, purified and stored at −80 °C.

### Vaccination

Two groups of 150 fish were tagged and i.p. injected with 0.05 mL of either the test vaccine or sterile PBS (control group). For this, fish were randomly transferred from their stock tank with a net into a bucket containing tank water at 12 ± 2 °C. Thereafter, groups of 2-5 fish were transferred at a time to a further bucket with MS-222 for anaesthesia and tagging. On a clean worktable each fish was marked using the Visible Implant Elastomer tagging system (VIE, Northwest marine technology, Inc). Mark colour was determined as orange for mock vaccinated and blue for the vaccinated fish (Figure 1).

**Figure 1.**
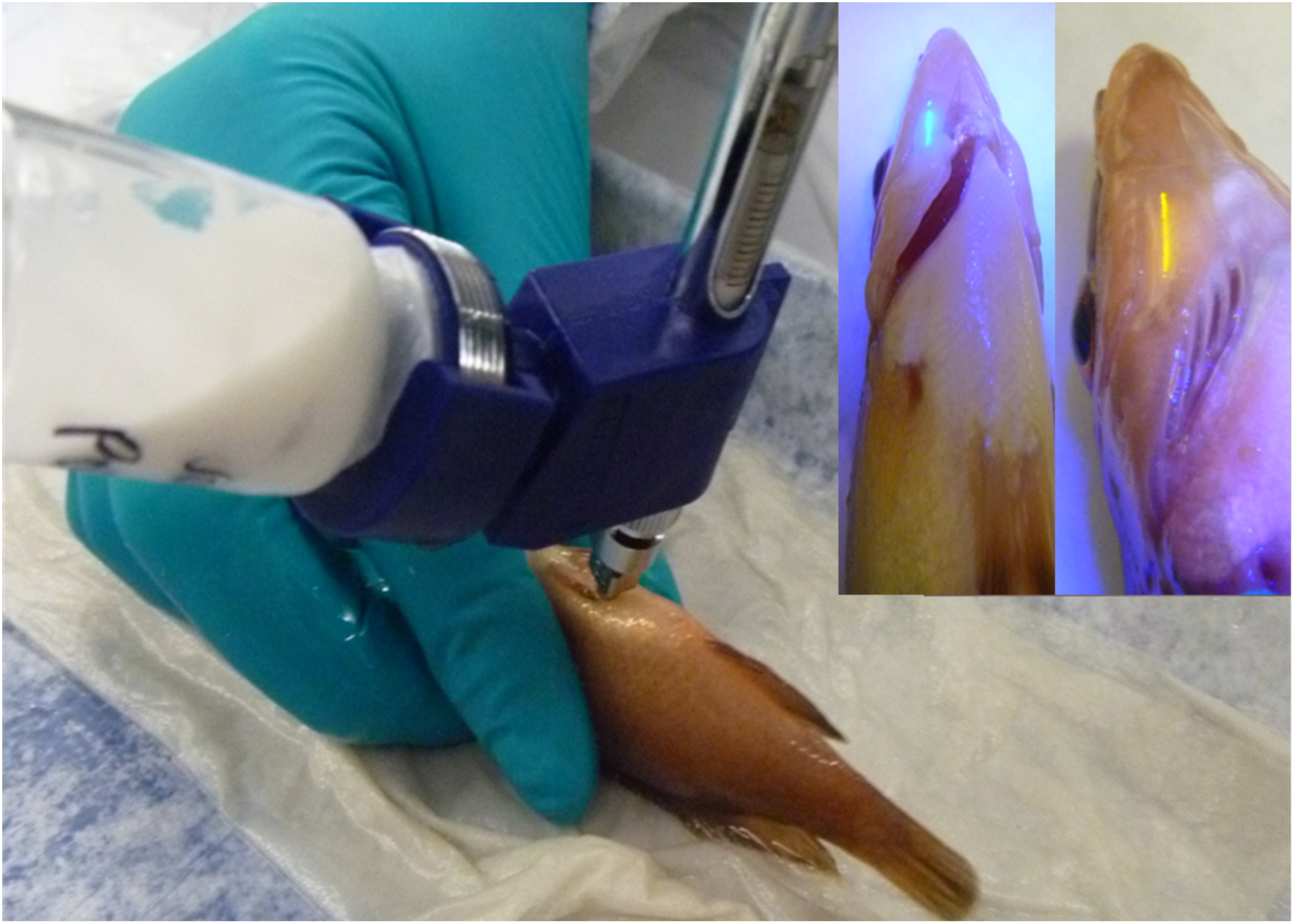
Intraperitoneal injection vaccination of ballan wrasse with oil-adjuvanted autogenous vaccine. Inset: wrasse tagged with Visible Implant Elastomer tagging system. Blue for the vaccinated fish and orange for mock vaccinated.

Immediately after tagging each fish was injected with the appropriate treatment using an automatic gun for the group of vaccinates and a sterile syringe for the mock vaccinates. For this, fish were i.p. vaccinated through the ventral wall of the coelomic cavity, one pelvic fin length anterior to the pelvic girdle and transferred directly into their holding tank at 12 ± 0.5 °C to recover (Figure 1). Vaccinated fish were then divided into 4 tanks (300 L with artificial plastic kelp and sections of plastic pipes to provide hides to the fish), 2 containing 75 fish vaccinated fish each and 2 tanks containing 75 mock vaccinated fish each.

Fish were held for 65 days at 12 ± 0.5 °C (780 DD) and blood samples collected from the caudal vein on days 31 and 65 post vaccination (prior to challenge) from 15 fish of each tank. The blood samples were centrifuged immediately after collection at 3,000 x g for 10 min and serum kept at −20 °C until used for serological analyses.

### Vaccine efficacy testing

After the immunisation period was completed, vaccinated and control fish were challenged with the two most virulent strains *i.e.* TW4/14 (a*As vapA* type V) and TW3/14 (a*As vapA* type V) using a tag and mix model with two different doses (pseudo replicate tanks), here referred as medium and high for isolate TW4/14 and high and very high for isolate TW3/14 as detailed in Table 3.

**Table 3.** Experimental design of the vaccine efficacy trial and relative percent survival (RPS) results.

| Species and Isolate | Tank and (n) | Dose type | cfu/fish | RPS (%) |
| --- | --- | --- | --- | --- |
| *aAs vapA* type V (TW4/14) | T03-10; 90 fish (45v +45c) | Medium | $1.0 \times 10^6$ | 95 |
| *aAs vapA* type V (TW4/14) | T03-09; 90 fish (45v +45c) | High | $1.0 \times 10^7$ | 91 |
| *aAs vapA* type V (TW3/14) | T03-08; 90 fish (45v +45c) | High | $1.0 \times 10^7$ | 79 |
| *aAs vapA* type V (TW3/14) | T03-07; 90 fish (45v +45c) | Very high | $1.0 \times 10^8$ | 20 |

V= vaccinated; C= control

### Infection and vaccination experiments: observations and sampling

For all the infection experiments, fish were observed at least twice a day. Diseased fish were classified as moribund or near moribund (humane endpoints) based on clinical signs (typically extreme lethargy when approached with a hand net). They were then euthanised by overdose of anaesthetic followed by confirmation of death by brain destruction, a UK Animals (Scientific Procedures) Act 1986 Amended Regulations (SI 2012/3039) Schedule 1 approved method (S1-M). All euthanised and dead fish were recorded throughout the experiments and accounted for posterior statistical analyses. To confirm specific mortalities, all moribund fish were necropsied, checked for gross pathology and sampled for bacteriology and histopathology as previously described. The challenge experiments were typically concluded when there was a period of at least five days with no mortalities. At the end of the vaccine efficacy trial, all surviving fish were killed by S1-M and blood sampled and processed as described before to measure specific antibody levels in the serum by ELISA.

All the experimental infections and vaccine efficacy tests were performed at 15 °C. Water temperatures were gradually increased over an acclimation period of 5-7 days prior to challenge.

### Specific IgM response

An indirect enzyme-linked immunosorbent assay (ELISA) was developed to detect and estimate the levels of specific anti-a*As* IgM in the ballan wrasse sera pre-vaccination and when the immunisation period was completed. Six samples including 2 replicates from each group were used for assessment of specific antibody responses by ELISA.

Antibody titres were determined according to the protocols outlined by [28] with modifications. Briefly, 96 – well ELISA plates (Immulon 4HBX, Thermo Scientific) were coated with 50 μL of 0.05% w/v poly – L– lysine in carbonate – bicarbonate buffer (0.05 M carbonate-bicarbonate pH 7.4, Sigma-Aldrich, St. Louis, UK) and incubated for 60 min at room temperature (RT). Plates were then washed 2 times with a low salt wash buffer (LSWB) (0.02 M Trizma base, 0.38 M NaCl, 0.05% Tween-20, pH 7.3). Bacteria i.e. a*As* type V isolate TW4/14, 100 μL at 10^8^ cfu/mL (OD_600_ 1.0), were then added to each well and plates were incubated overnight at 4 °C. The bacteria were previously prepared by growing them on TSB at 22 °C for 48 h with continuous shaking at 150 rpm and washed 2 times with PBS, resuspended and adjusted to an OD_600_ 1.0 prior to 96 – well plates inoculation. Glutaraldehyde (50 μL, 0.05% (v/v)) diluted in PBS was added to the wells of the ELISA plate to fix the antigen, incubated 20 min at RT and plates were washed 3 times with LSWB.

The plates were then post-coated with 3% w/v casein in distilled water (250 μL) to block non-specific binding sites and incubated for 180 min at RT. The supernatant was decanted and plates were stored at −20 °C for up to 3 weeks. LSWB was used to wash the plate 3 times and 100 μL of hydrogen peroxide (H_2_O_2_; 0.3% of 10% stock solution in 10% methanol) was added to each well to quench endogenous peroxidase activity of the bacteria and incubated for 30 min at RT.

Diluted serum (100 μL per well; from 1:50 to 1:800) in 0.5% casein (w/v) and in PBS, were added to the plates and incubated for 1 h at RT. Plates were washed 5 times with high salt wash buffer (HSWB) (0.02 M Trizma base, 0.5 M NaCl, 0.1% Tween-20, pH 7.4) and were incubated with the last HSWB wash for 5 min at RT.

Anti – Asian sea bass IgM MAbs (ADL, Stirling, UK) (shown to cross react with ballan wrasse IgM) diluted 1/50 with 0.01% Bovine Serum Albumin (BSA) in PBS was then added to each well (100 μL), and incubated for 1 h at RT. The plates were then washed 5 times in HSWB as described above. Goat anti – mouse – horseradish peroxidase (HRP) conjugate (Sigma-Aldrich, UK) diluted 1/4000 with 0.01% BSA in LSWB was then added to the plates. Chromogen in substrate buffer (prepared by adding 150 μL of chromogen 42 mM trimethyl-benzidine, TMB to 15 mL of substrate buffer containing 5 μL H_2_O_2_ in 6 mL of 50% acetic acid) was then added (100 μL / well) for assay development.

The plates were incubated for 3 – 5 min at RT and the reaction stopped by adding 50μL sulfuric acid (2 M H_2_SO_4_). The absorbance was measured at OD_450_ using a 96 – well plate spectrophotometer (Biotek Instruments, Friedrichshall, Germany). The sensitivity threshold of the assay was determined as 3x the absorbance value of wells containing PBS (background absorbance). Samples above this value were considered positive for specific antibodies.

### Statistical analyses

The efficacy/potency of the vaccine was assessed by calculating the relative percent survival (RPS) which indicates the proportional percentage between the cumulative (cm) morbidities of vaccinated group and cumulative morbidities of mock vaccinated group using the equation below [29].

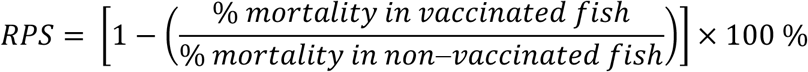

Minitab 18 was used to produce Kaplan – Meier survival curves and perform log-rank non-parametric tests (significance level p < 0.05) for survival comparisons. Antibody responses in serum samples of vaccinated and non – vaccinated ballan wrasse were tested for normality (Anderson-Darling test) and homogeneity of variance (Levene’s test). Kruskal-Wallis non – parametric test was used for dose response assessment in relation to antibody titres while a pairwise comparison (Mann Whitney-U test (CI = 95%) was conducted between the antibody responses.

### Ethical statement

Bacterial infection and vaccination procedures were performed under the authority of UK Government Home Office project licences, following approval by the Animal Welfare and Ethical Review Body (AWERB) at the Centre for Environment, Fisheries and Aquaculture Science (Cefas) and University of Stirling. Ballan wrasse were treated in accordance with the Animals (Scientific Procedures) Act 1986 Amended Regulations (SI 2012/3039).

## RESULTS

### Infection experiments

In the first two infection experiments, injection with high doses of a*As* type V isolates TW3/14, TW4/14 and TW187/14 and type VI isolate TW164/15 resulted in 100% moribundity/mortality by 7 days post challenge (Table 2). Clinical signs were first recorded at 4 days post infection (dpi) for both *vapA* types and 100% morbidities achieved by 4 and 8 dpi, for *vapA* types V and VI respectively. In all experiments, the a*As* isolates were recovered from moribund fish as pure cultures (punctate whitish to greyish colonies) from swabbed internal organs. The a*As* type VI isolate tested produced a diffusible pigment, (brown on TSA and grey on BA) that became more evident after five days incubation (Supplementary File1). A representative isolate from each strain was stored at −80 °C under Cefas bacterial culture collection codes 17032, 17033 and 17034 after being in vivo passaged in fish.

The infections performed with *Aliivibrio salmonicida*, *Allivibrio logei*, *Vibrio splendidus* and *Vibrio ichtyoenteri* did not cause any sign of disease or mortalities after 7 dpi in infection experiment 1 (Table 2). In infection experiment 2, only *Aliivibrio salmonicida* caused 2 mortalities (13%) on day 3 and the other three were not pathogenic.

In infection experiment 3 (Table 2), fish infected with a medium dose of a*As vapA* type VI isolates TW184/16 (1.6×10^7^ cfu/fish) and TW164/15 (3×10^7^ cfu/fish) resulted in mortalities of 25% and 33% respectively, while a high dose (10^9^ cfu/fish) caused 100% mortalities for both isolates. The two a*As* type VI isolates presented brown pigment as described before. Signs of disease presented more rapidly for isolate TW184/16 for medium (6 dpi) and high (2 dpi) dose but similar to those of TW164/15 (7 and 3 dpi, respectively) (Supplementary File 2). Morbid fish showed some signs of reduced appetite often followed by imbalance, lethargy and full loss of equilibrium. Gross external pathology included ascites, and occasionally haemorrhaging at the injection site, internally liquefaction of organs and white deposits in the peritoneum (Supplementary File 3). More liquefaction was noted with TW184/16 than TW164/16. Interestingly the bacterium was not isolated from any survivor fish challenged with medium dose at termination on 14 (TW184/16) and 16 (TW164/15) dpi and there were no any obvious external or internal signs of disease in them.

The *Photobacterium indicum* isolates TW138/16 and TW181/16 both caused 11 (92%) overnight mortalities when administered at high doses of 3.2×10^9^ and 9×10^9^ cfu/fish respectively. The remaining fish injected with a high dose of TW138/16 was removed on day 3 post infection while the last fish injected with TW181/16 were euthanised on welfare grounds at day 6 post infection (Supplementary File 4). For the *Photobacterium indicum* challenges with medium doses (3.2×10^7^ and 9×10^7^ cfu/fish), of isolates TW181/16 and TW138/16 resulted in 8/12 and 6/12 mortalities respectively by day 3 post challenge (Supplementary File 4). The remaining fish (n= 4) in the tank challenged with TW181/16 were terminated at day 9 post infection as no morbidities occurred for 3 days and all presented lesions at the injection site during the daily observations. In the tank infected with medium dose of isolate TW138/16 a single dead fish with a large lesion around the injection site was removed on day 9, while monitoring of the remaining fish (n= 6) continued. On day 16 post infection all surviving fish (n=6) presented severe ventral lesions at the i.p. injection site (Supplementary File 5) and some of these lesions extended into the cavity and for this reason the experiment was terminated. Morbid fish infected with *Photobacterium indicum* showed a reduced feeding response often followed by imbalance, lethargy and full loss of equilibrium with a very rapid progression (< 24 h) of the signs. During the necropsies, the majority of the fish had ascites, liquefaction of organs and swelling coelomic cavity due to ascites. Internally, haemorrhages or lesions were noted and the severity of these progressed over time.

Additional testing of isolate a*As* type V (TW4/14) in infection experiments 4 and 5 confirmed this organism was virulent. Morbidities were recorded within 2 dpi with the high dose (10^8^ cfu/fish) and 100% mortalities were reached by day 4. A dose of 10^7^ cfu/fish reached a maximum of 87% mortality by day 6 post infection, while no signs of disease or morbidities were noted for fish challenged with the lowest dose (10^4^ cfu/fish) (Supplementary File 6). Similar results were obtained in the second pre-test, with 53% for group exposed to 10^6^ cfu/fish and only 7% in the group exposed to 10^4^ cfu/fish. (Supplementary File 7). The predicted LD_60_ based on these experiments was between 10^6^ cfu/fish (53%) and 10^7^ cfu/fish (87%), with 10^7^ cfu/fish selected as one of the doses for the vaccine efficacy trials.

For the additional testing undertaken with a*As* type VI (TW164/15) in infection experiment 4, only a single morbidity occurred at 3 dpi, while the rest of the fish showed no visual signs of disease or adverse behaviour. The trial was terminated at 20 dpi and fish (n= 6) sampled for bacteriology. All inoculated plates were considered negative as no significant bacterial colonies were observed. For these reasons this isolate was not used for the vaccine testing and a replacement isolates was selected as described below.

### Virulence determination

LD_50_ values for the different isolates by day 8 post infection were calculated based on results from all the infection experiments. Atypical *Aeromonas salmonicida vapA* type V isolate TW4/14 was the most virulent followed by a*As vapA* type V isolate TW3/14, *Photobacterium indicum* and a*As vapA* type VI. The a*As vapA* type V isolates (TW3/14, TW4/14 and TW187/14) were chosen for macrorestriction analysis using PFGE to select a replacement isolate for a*As vapA* type VI (TW164/15) which was not virulent during experiment 3. Differences were observed in the restriction sites for isolate TW3/14 in comparison to TW4/14 and TW187/14 (Figure 2) which may explain the differences in virulence mentioned above. The *Aliivibrio logei*, *Vibrio splendidus*, *Aliivibrio salmonicida* and *Vibrio ichtyoenteri* were not pathogenic. The average of the 3 estimations for a*As vapA* type V (TW4/14) was 3.6×10^6^ cfu/fish. The average of the 2 estimations for *Photobacterium indicum* was 2.2×10^7^ cfu/fish. A comparison of all the LD_50_ values of the different isolates is presented in Table 4.

**Table 4.** Lethal dose 50% (LD_50_) of 6 bacterial species used in the trials by day 8 post infection.

| Bacterial species | Isolate | LD<sub>50</sub> (cfu/fish) |
| --- | --- | --- |
| Atypical *Aeromonas salmonicida* type V | TW4/14* | 2.0 x 10<sup>5</sup> |
| Atypical *Aeromonas salmonicida* type V | TW4/14 | 2.8 x 10<sup>6</sup> |
| Atypical *Aeromonas salmonicida* type V | TW4/14** | 6.1 x 10<sup>6</sup> |
| Atypical *Aeromonas salmonicida* type V | TW3/14** | 1.6 x 10<sup>7</sup> |
| *Photobacterium indicum* | TW181/16 | <3.2 x 10<sup>7</sup> |
| *Photobacterium indicum* | TW138/16 | 1.3 x 10<sup>8</sup> |
| Atypical *Aeromonas salmonicida* type VI | TW184/16 | 3.4 x 10<sup>8</sup> |
| Atypical *Aeromonas salmonicida* type VI | TW164/15 | 5.3 x 10<sup>8</sup> |
| *Aliivibrio logei* | TW242/16 | >1.5 x 10<sup>9</sup> |
| *Aliivibrio logei* | TW186/16 | >1.8 x 10<sup>9</sup> |
| *Vibrio splendidus* | TW130/16 | >3.0 x 10<sup>9</sup> |
| *Aliivibrio salmonicida* | TW322/16 | >5.0 x 10<sup>9</sup> |
| *Vibrio ichthyoenteri* | TW319/16 | >8.0 x 10<sup>9</sup> |

(\*) passaged isolate used in dose trials; (\*\*) data from mock vaccinated.

**Figure 2.**
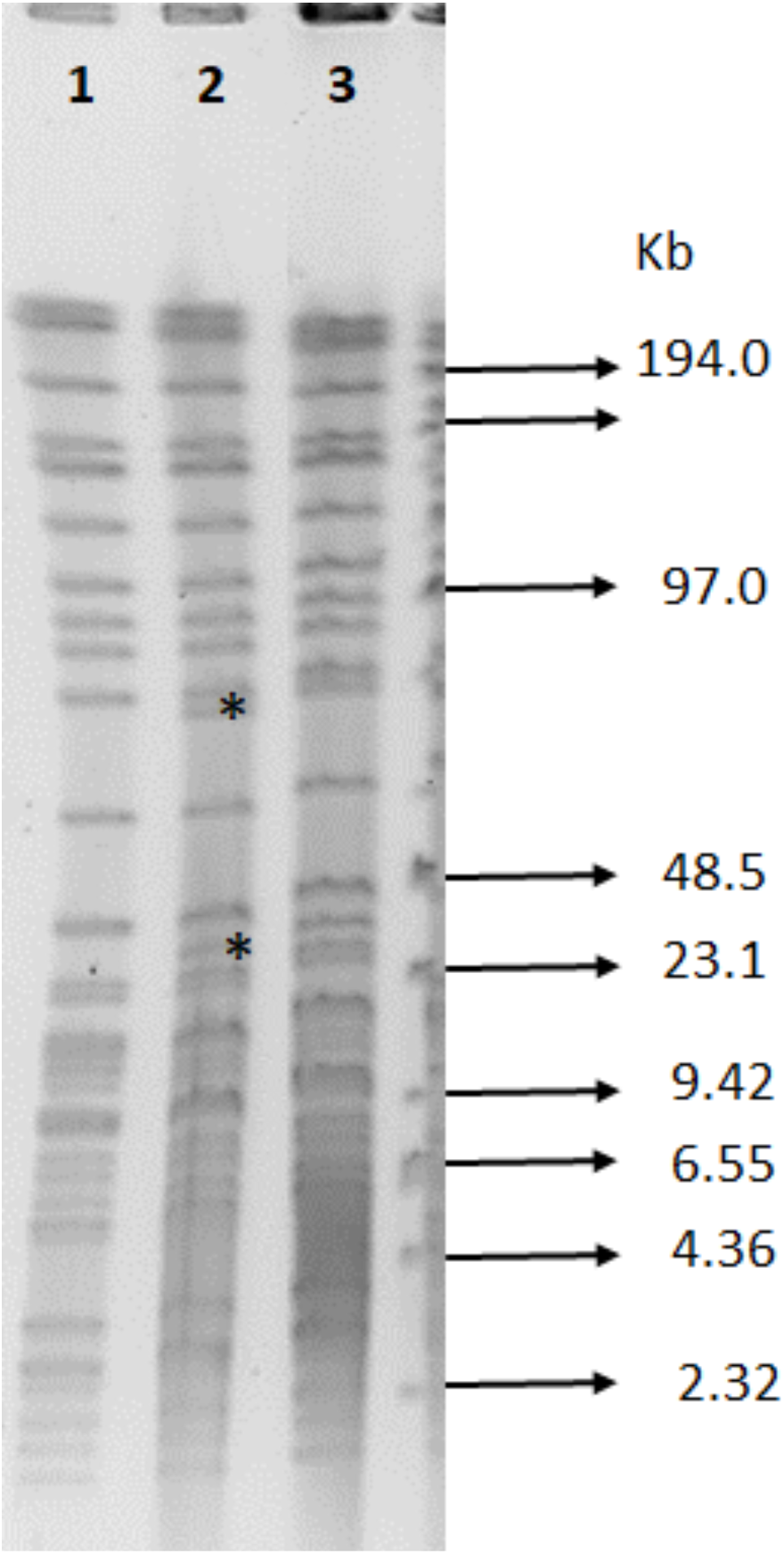
Pulsed-filed gel electrophoresis patterns of *Aeromonas salmonicida vapA* type V isolates TW4/14, TW 187/14 and TW 3/14 restricted with *SpeI* enzyme (New England Labs, UK). From left to right, TW 3/14 (position 1), TW 4/14 (position 2) and TW 187/14 (position 3). Molecular marker mixture of lambda DNA-Hind III fragments and lambda concatamer; 48±5 kb (Low Range PFG Marker, New England Labs, UK). Notice the difference between pulsotype profiles for isolates TW 3/14 and TW 4/14 and TW 187/14 (asterisk).

### Vaccine efficacy

Significant protection was demonstrated with vaccinated fish experiencing significantly lower mortalities than control fish when challenged with a*As* type V from isolates TW4/14 and TW3/14. First morbidities were recorded at 5 dpi (TW3/14) and 6 dpi (TW4/14) in the mock-vaccinated groups, and 7 dpi (TW3/14) and 15 dpi (TW4/14) in the vaccinated groups (Figure 3A, B).

**Figure 3.**
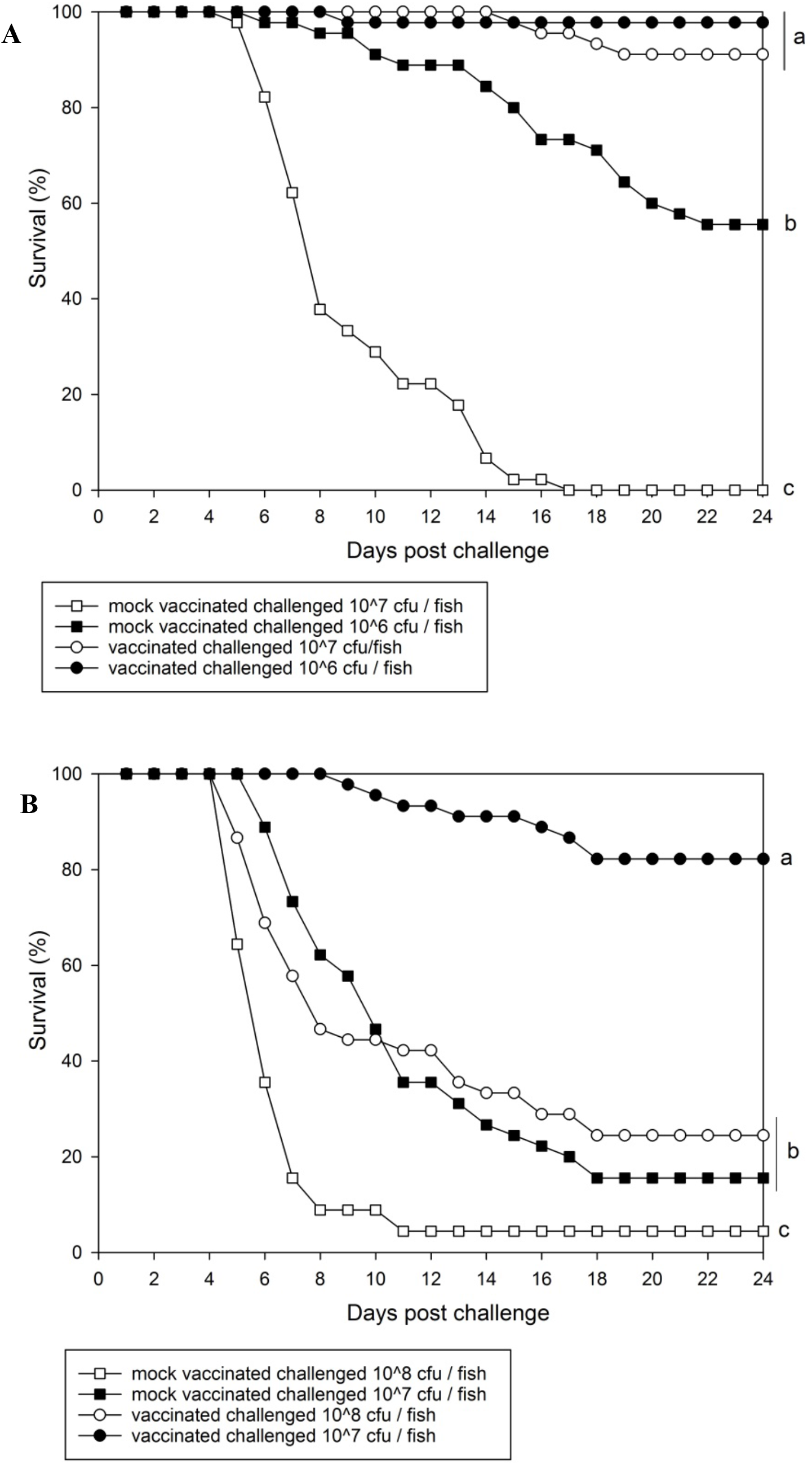
Survival (%) of i.p. injected vaccinated and non – vaccinated ballan wrasse challenged with A) a*As* type V (isolate TW3/14) at high and very high doses and B) a*As* type V (isolate TW4/14) at medium and high doses. Letters represents statistical significance (p < 0.05).

Mortalities were significantly higher for mock-vaccinated fish challenged with either of the two isolates over a period of 24 days (Figure 3A, B). Isolate TW3/14 at a very high dose of 1×10^8^ cfu/fish resulted in 96% mortality relative to the control fish and 34% mortality relative to the vaccinated group. With the same isolate, a dose of 10^7^ cfu/fish caused 84% mortality relative to the control groups and only 18% relative to the vaccinated group. Isolate TW4/14, at a high dose of 10^7^ cfu/fish, caused 100% mortalities relative to the control and only 9% relative to the vaccinated group while a medium dose of 10^6^ cfu/fish caused 44% in the control groups and only 2% to the vaccinated fish.

The RPS values in the ballan wrasse challenged with isolate TW4/14 at medium and high doses were 95% (10^6^ cfu/fish) and 91% (10^7^ cfu/fish), respectively. The group exposed to strain TW3/14 had an RPS of 79% with the high dose of 10^7^ cfu/fish but a low RPS (20%) was recorded for the group injected with the very high challenge dose (10^8^ cfu/fish) (Table 4). Despite the low RPS, the survival of vaccinated fish in this group was still significantly higher when compared with the mock vaccinated group (Figure 3B).

### Specific IgM response

Non – specific binding was observed in the preliminary results (Supplementary File 8). This was reduced when plates were treated with 0.3% hydrogen peroxide to quench endogenous peroxidase activity of the bacteria and when 0.5% casein and 0.01% BSA were included in the fish serum and Anti – Asian sea bass IgM MAbs, respectively.

A very weak antibody response was noted for serum samples collected from mock-vaccinated fish (controls) prior to challenge and these were considered negative. The vaccinated fish had significantly higher mean antibody titres at all sera dilutions in contrast to mock-vaccinated fish (Figure 4).

**Figure 5.**
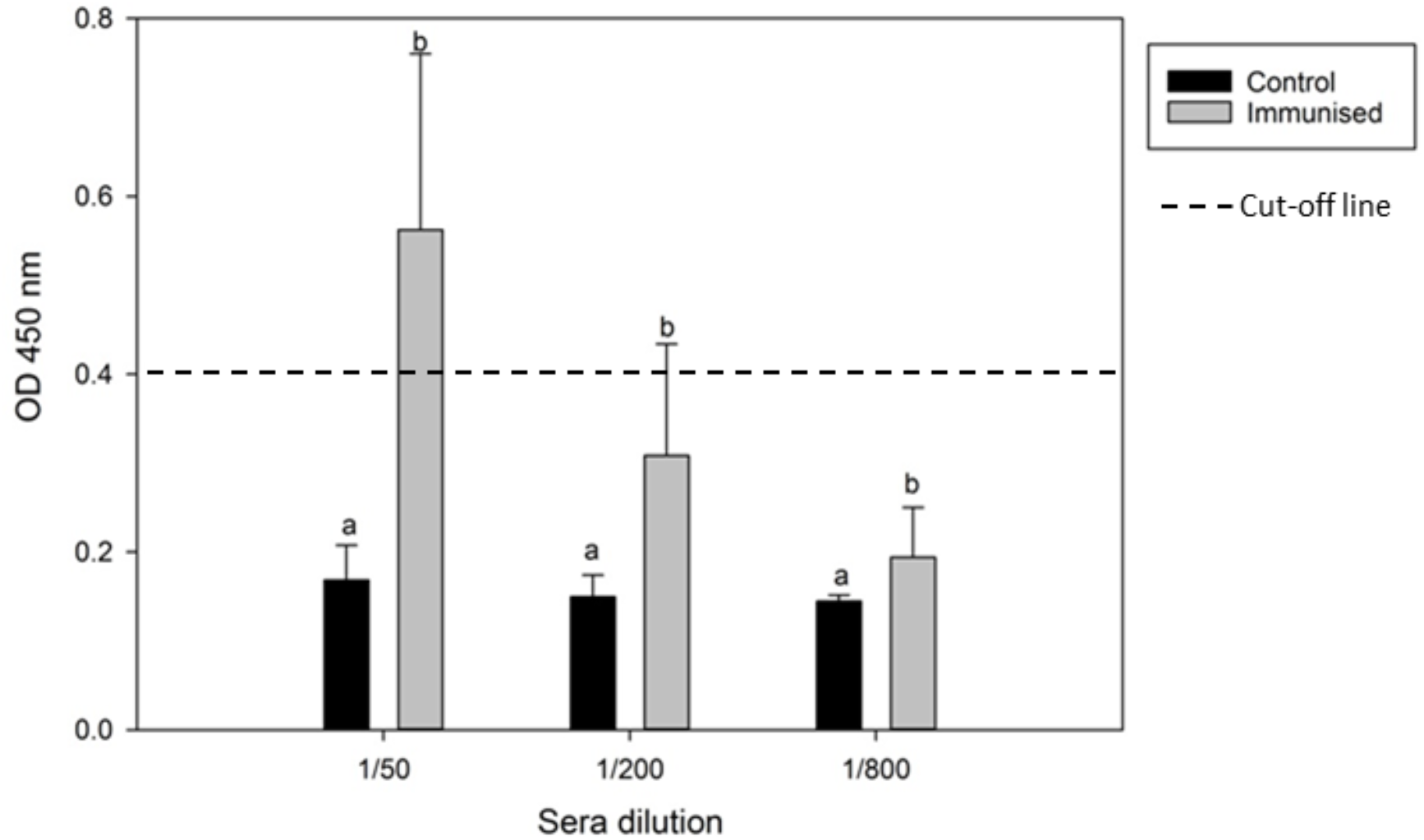
Ballan wrasse specific antibody (IgM) response to a*As* pre-vaccination (control, n= 6 samples x 2 replicates) and after immunisation period was completed (780 DD after i.p. vaccination, n= 6 replicates x 2 replicates). Letters represents statistical significance (p < 0.05).

## DISCUSSION

In the present study, the virulence of a*As* type V and VI, *Aliivibrio logei*, *Aliivibrio salmonicida*, *Vibrio splendidus*, *Vibrio ichthyoenteri* and *Photobacterium indicum* were assessed. The results obtained confirmed that a*As vapA* type V was the most pathogenic of all the bacterial species (followed by *Photobacterium indicum*, a*As* VI and the rest of the *Vibrionaceae*). Importantly, the vaccine tested was highly protective against a*As* type V and significantly higher titres of specific systemic IgM were detected in vaccinated fish when compared to controls.

The virulence studies confirmed that a*As vapA* type V (from both isolates tested) were highly virulent in ballan wrasse when i.p. injected. The RPS obtained with medium and high doses for *vapA* type V from isolate TW4/14 (95% and 91%, respectively) and high dose of *vapA* type V from isolate TW3/14 (79%) strains, confirmed the effectiveness of the injection vaccine against homologous strains of a*As* and were in agreement with previous results conforming that ballan wrasse can be effectively immunised by i.p. injection against this pathogen [17]. When vaccinated ballan wrasse were challenged with a very high dose (10^8^ cfu/fish) of the strain TW3/14 RPS was only of 20% suggesting that high challenge dose may have suppressed or overwhelmed protective memory responses. This highlights the relevance of biosecurity and good farming practices to maintain the pathogens challenge pressure as low as possible during the production cycle.

A specific antibody response (IgM) to the vaccine was measured in immunised fish at 780 DD which was significantly higher compared to non-vaccinated fish. The high RPS levels in vaccinated fish and specific antibody response following vaccination are indicators that the vaccine indeed triggered a specific protective humoral response against a*As*. Similar responses have been induced in other species immunised with typical or atypical stains of *As*, such as lumpsucker [30, 31], Atlantic salmon [32, 33], rainbow trout [34] and spotted wolfish [35]. A high quantity of cytoplasmic peroxidases (*e.g.* thiol peroxidase) have previously been reported in *A. salmonicida* cells [36] and high antigen endogenous peroxidase activity appeared to cause substantial background during ELISA development. This background was quenched using hydrogen peroxide prior to antibody-antigen complexing. However, a high absorbance threshold of the ELISA could not be avoided using our cut-off criteria (3x background OD = 0.4), thus a 1/50 test titre was the most preferable to use to determine positive antibody responses to a*As* vaccination. Nonetheless, the titre of antibodies was consistently higher in vaccinated fish up to and including a dilution of 1/800. These results suggested that antibodies might be involved in protection against a*As*.

Interestingly, differences in virulence were observed for two atypical *Aeromonas salmonicida vapA* type V isolates (TW3/14 and TW4/14), with the latter being the most virulent. Microrestriction and PFGE analysis corroborated these results, indicating small but potentially important genomic differences between isolates. Atypical *Aeromonas salmonicida* isolates heterogeneity has been previously assessed with the same method [26]. Characterisation of all the available a*As vapA* types for ballan wrasse by PFGE will be beneficial to select isolates for future vaccine formulations.

The *vapA* VI isolates appeared less virulent than a*As* type V and similar results were reported for Norwegian a*As* type V and VI isolates [17]. In that study a*As* type V induced high mortalities (75 – 89% morbidities) in 50 g ballan wrasse when i.p. injected with 10^7^ cfu /fish and also by cohabitation (51%). The type VI isolates were less virulent, in particular by cohabitation (8%) than i.p. injection (70 – 85%). Interestingly, in the present study, survivor fish infected with a*As* type VI at a medium dose (10^7^ cfu / fish) did not show any obvious external or internal signs of the disease and no bacteria were recovered from those fish suggesting that ballan wrasse were able to clear the infection. This is in agreement with a previous study that reported similar responses in survivors from groups infected with a*As* type V and VI [17].

The *Aliivibrio logei*, *Vibrio splendidus* and *Vibrio ichthyoenteri* isolates were not pathogenic to ballan wrasse by i.p. injection even when very high challenge doses were administered. *Aliivibrio salmonicida* was the only pathogen that caused mortalities (13%) but only when very high infection dose of 5 × 10^9^ cfu/fish was administered.

The a*As vapA* type VI isolates occasionally displayed a peculiar alternative morphology that included the presence of large greyish and small transparent colonies (Supplementary File 9). Previous reports have documented this phenomenon associated with variable expression of a functional A-layer and consequently with variable virulence [22, 37–41]. Although in the present study, the inclusion of isolates displaying such alternative morphology was generally avoided, this should not be ruled out as a possible explanation behind the lack of virulence observed in this experiment.

In previous reports of experimental infections with a*As* type VI isolates in cleaner fish, lumpsucker succumbed to disease at lower doses of 2 ×10^3^ cfu/mL (bath) and 4 × 10^4^ cfu /fish (i.p. injection) after exposure [18]. Other fish species like spotted wolfish also experienced high mortalities with low doses of a*As* (10^3^ and 10^4^ cfu / mL) by i.p. injection [8, 15]. In contrast, turbot required very high doses (10^8^ and 10^10^ cfu / mL) for mortalities to be induced by i.p. injection [16]. As suggested previously, there is a strong association between host species and *vapA* type and it is possible that *vapA* type V is more strongly associated with wrasse than lumpfish and vice versa for type IV [42].

This study is the first experimental confirmation that *Photobacterium indicum* can be pathogenic towards wrasse, through fulfilment of Koch’s postulates. *Photobacterium indicum* was regularly isolated from diseased ballan wrasse during disease surveys in Scotland and it was linked to histopathological lesions [4]. There are no previous reports on fish susceptibility to *Photobacterium indicum* although it has been isolated from moribund American lobster (*Homarus americanus*) associated with stress and has been reported as an opportunistic pathogen for this crustacean species [43, 44]. In cleaner fish, *Photobacterium* sp. was recently recovered from lumpsucker experiencing mortalities due to *Pseudomonas anguilliseptica* under rearing conditions in Scotland [45]. The pathogenicity results for *Photobacterium indicum* obtained in the present study need to be treated with caution as disease was induced only via i.p. injection which bypasses the natural protective mucosal barriers of the host *e.g.* skin, gills and gastrointestinal tract [46, 47]. Signs of disease and gross pathology for *Photobacterium indicum* were similar to those seen with a*As* with moribund fish showing reduced feeding response often followed by imbalance, lethargy and full loss of equilibrium. The peritoneal cavity of the diseased fish was extended (ascites) and internally, liquefaction was observed in all the organs. Granulomatous formations were seen in livers of moribund fish infected with a*As* which concurred with previous reports [17]. This needs to be considered when performing differential diagnosis based on gross pathology and clinical signs. Ventral lesions at the i.p. injection site were observed on survivor fish, which may be related to localised immune responses at the injection site [48, 49].

## CONCLUSIONS

This study developed an i.p. injection challenge model was for ballan wrasse against Scottish a*As vapA* type V isolates and this was used to test the efficacy of an injectable autogenous multivalent vaccine. The vaccination results obtained in the current study are very encouraging as they confirmed i.p vaccination can be used as means to control and potentially eliminate morbidities in ballan wrasse hatcheries and cage sites due to a*As vapA* type V and likely other *vapA* types. The vaccine tested was highly protective against medium and high challenge doses of a*As* type V from two different isolates (RPS 79-95%) and significantly higher titres of specific systemic IgM were detected in vaccinated fish when compared to controls. In addition, this study provided important new data on the pathogenicity and virulence of routinely recovered bacterial species from diseased ballan wrasse. The pathogenicity and virulence of *Photobacterium indicum* to ballan wrasse is reported for the first time. Atypical *Aeromonas salmonicida* type V isolate TW4/14 was the most virulent pathogen followed by a*As* type V isolate TW3/14, *Photobacterium indicum* and atypical a*As* type VI (including isolate from lumpsucker). The *Vibrio* spp. and *Aliivibrio* spp. were not pathogenic by i.p. injection to the ballan wrasse population tested. Further work is needed to assess the efficacy of vaccination against other isolates that have proven to be pathogenic such as a*As* type VI and *Photobacterium indicum* and to explore the feasibility of immersion vaccination strategies as the species encounters the pathogens at earlier life stage (< 25 g) and this immunisation route is desirable for juvenile ballan wrasse in the hatcheries. In addition, full characterisation should be performed on a*As* isolates within the same *vapA* types.

## ACKNOWLEDGEMENTS

The authors would like to thank Otterferry Seafish Ltd. for the provision of the fish and the aquarists and bacteriology team at Cefas for their technical assistance during the trial.

## REFERENCES

[1] B. Austin, D.A. Austin, Aeromonadaceae representative (Aeromonas salmonicida), Bacterial fish pathogens, Springer 2016, pp. 215–321.

[2] A.J. Brooker, A. Papadopoulou, C. Gutierrez, S. Rey, A. Davie, H. Migaud, Sustainable production and use of cleaner fish for the biological control of sea lice: recent advances and current challenges, Veterinary Record 183(12) (2018) 383.

[3] B. Hjeltnes, B. Bang-Jensen, G. Bornø, A. Haukaas, C.S. Walde, The Health Situation in Norwegian Aquaculture 2018, Norwegian Veterinary Institute (2019) 132.

[4] A. Papadopoulou, T.S. Wallis, J.G. Ramirez-Paredes, S.J. Monaghan, A. Davie, H. Migaud, A. Adams, Atypical Aeromonas salmonicida vapA type V and Vibrio spp. are predominant bacteria recovered from ballan wrasse Labrus bergylta in Scotland, Diseases of aquatic organisms In press (2020).

[5] P.J. Midtlyng, Vaccination against Furunculosis, in: R. Gudding, A. Lillehaug, Ø. Evensen (Eds.), Fish Vaccination, John Wiley & Sons, Ltd, Chichester, 2014, pp. 185–199.

[6] B. Björnsdóttir, S. Gudmundsdóttir, S. Bambir, B. Gudmundsdóttir, Experimental infection of turbot, Scophthalmus maximus (L.), by Aeromonas salmonicida subsp. achromogenes and evaluation of cross protection induced by a furunculosis vaccine, Journal of Fish Diseases 28(3) (2005) 181–188.

[7] V. Lund, L.M. Jenssen, M.S. Wesmajervi, Assessment of genetic variability and relatedness among atypical Aeromonas salmonicida from marine fishes, using AFLP-fingerprinting, Diseases of aquatic organisms 50(2) (2002) 119–126.

[8] V. Lund, H. Mikkelsen, M.B. Schrøder, Comparison of atypical furunculosis vaccines in spotted wolffish (Anarhicas minor O.) and Atlantic halibut (Hippoglossus hippoglossus L.), Vaccine 26(23) (2008) 2833–2840.

[9] P. Pylkkö, Atypical Aeromonas salmonicida-infection as a threat to farming of arctic charr (Salvelinus alpinus L.) and european grayling (Thymallus thymallus L.) and putative means to prevent the infection, University of Jyväskylä2004.

[10] Y. Attia, I. Schmerold, A. Hönel, The legal foundation of the production and use of herd-specific vaccines in Europe, Vaccine 31(36) (2013) 3651–3655.

[11] E.M.A. CMDv, Recommendations for the manufacture, control and use of nactivated autogenous veterinary vaccines within the EEA, London 2017, p. 14.

[12] S.R. Haskell, K. Carberry-Goh, M.A. Payne, S.A. Smith, Current status of aquatic species biologics, J Am Vet Med Assoc 225(10) (2004) 1541–4.

[13] M. Saléry, Autogenous vaccines in Europe: national approaches to authorisation, (2017).

[14] B.K. Gudmundsdóttir, B. Björnsdóttir, Vaccination against atypical furunculosis and winter ulcer disease of fish, Vaccine 25(30) (2007) 5512–5523.

[15] M. Ingilæ, J.A. Arnesen, V. Lund, G. Eggset, Vaccination of Atlantic halibut Hippoglossus hippoglossus L., and spotted wolffish Anarhichas minor L., against atypical Aeromonas salmonicida, Aquaculture 183(1) (2000) 31–44.

[16] Y. Santos, S. García-Marquez, P. Pereira, F. Pazos, A. Riaza, R. Silva, A. El Morabit, F. Ubeira, Efficacy of furunculosis vaccines in turbot, Scophthalmus maximus (L.): evaluation of immersion, oral and injection delivery, Journal of fish diseases 28(3) (2005) 165–172.

[17] E. Biering, Ø. Vaagnes, B. Krossøy, S. Gulla, D. Colquhoun, Challenge models for atypical Aeromonas salmonicida and Vibrio anguillarum in farmed Ballan wrasse (Labrus bergylta) and preliminary testing of a trial vaccine against atypical Aeromonas salmonicida, Journal of fish diseases 39(10) (2016) 1257–1261.

[18] A. Rønneseth, G.T. Haugland, D.J. Colquhoun, E. Brudal, H.I. Wergeland, Protection and antibody reactivity following vaccination of lumpfish (Cyclopterus lumpus L.) against atypical Aeromonas salmonicida, Fish Shellfish Immunol 64 (2017) 383–391.

[19] B.K. Gudmundsdóttir, S. Gudmundsdóttir, Evaluation of cross protection by vaccines against atypical and typical furunculosis in Atlantic salmon, Salmo salar L, Journal of Fish Diseases 20(5) (1997) 343–350.

[20] A. Klindworth, E. Pruesse, T. Schweer, J. Peplies, C. Quast, M. Horn, F.O. Glöckner, Evaluation of general 16S ribosomal RNA gene PCR primers for classical and next-generation sequencing-based diversity studies, Nucleic Acids Research 41(1) (2013) e1–e1.

[21] S. Yamamoto, H. Kasai, D.L. Arnold, R.W. Jackson, A. Vivian, S. Harayama, Phylogeny of the genus Pseudomonas: intrageneric structure reconstructed from the nucleotide sequences of gyrB and rpoD genesThe GenBank accession numbers for the sequences determined in this work are: gyrB, D37926, D37297, D86005–D86019 and AB039381–AB039492; rpoD, D86020–D86036 and AB039493–AB039624, Microbiology 146(10) (2000) 2385–2394.

[22] S. Gulla, V. Lund, A. Kristoffersen, H. Sørum, D. Colquhoun, vapA (A-layer) typing differentiates Aeromonas salmonicida subspecies and identifies a number of previously undescribed subtypes, Journal of fish diseases 39(3) (2016a) 329–342.

[23] N.D. Young, I. Dyková, B.F. Nowak, R.N. Morrison, Development of a diagnostic PCR to detect Neoparamoeba perurans, agent of amoebic gill disease, Journal of Fish Diseases 31(4) (2008) 285–295.

[24] OIE, Manual of diagnostic tests for aquatic animals, Chapter 2.3, (2018).

[25] K. Bartie, F.W. Austin, A. Diab, C. Dickson, T.T. Dung, M. Giacomini, M. Crumlish, Intraspecific diversity of Edwardsiella ictaluri isolates from diseased freshwater catfish, Pangasianodon hypophthalmus (Sauvage), cultured in the Mekong Delta, Vietnam, Journal of fish diseases 35(9) (2012) 671–682.

[26] M.-L. Hänninen, V. Hirvelä-Koski, Genetic diversity of atypical Aeromonas salmonicida studied by pulsed-field gel electrophoresis, Epidemiology & Infection 123(2) (1999) 299–307.

[27] L.J. Reed, H. Muench, A simple method of estimating fifty per cent endpoints, American Journal of Epidemiology 27(3) (1938) 493–497.

[28] A. Adams, K.D. Thompson, D. Morris, C. Farias, S.C. Chen, Development and use of monoclonal antibody probes for immunohistochemistry, ELISA and IFAT to detect bacterial and parasitic fish pathogens, Fish & Shellfish Immunology 5(8) (1995) 537–547.

[29] D.F. Amend, Potency testing of fish vaccines, Fish biologics: serodiagnostics and vaccines (1981) 447–454.

[30] T. Erkinharju, M.R. Lundberg, E. Isdal, I. Hordvik, R.A. Dalmo, T. Seternes, Studies on the antibody response and side effects after intramuscular and intraperitoneal injection of Atlantic lumpfish (Cyclopterus lumpus L.) with different oil-based vaccines, Journal of Fish Diseases 40(12) (2017) 1805–1813.

[31] A. Rønneseth, D.B. Ghebretnsae, H.I. Wergeland, G.T. Haugland, Functional characterization of IgM+ B cells and adaptive immunity in lumpfish (Cyclopterus lumpus L.), Developmental & Comparative Immunology 52(2) (2015) 132–143.

[32] A.B. Romstad, L.J. Reitan, P. Midtlyng, K. Gravningen, Ø. Evensen, Development of an antibody ELISA for potency testing of furunculosis (Aeromonas salmonicida subsp salmonicida) vaccines in Atlantic salmon (Salmo salar L), Biologicals 40(1) (2012) 67–71.

[33] A.B. Romstad, L.J. Reitan, P. Midtlyng, K. Gravningen, Ø. Evensen, Antibody responses correlate with antigen dose and in vivo protection for oil-adjuvanted, experimental furunculosis (Aeromonas salmonicida subsp. salmonicida) vaccines in Atlantic salmon (Salmo salar L.) and can be used for batch potency testing of vaccines, Vaccine 31(5) (2013) 791–796.

[34] K.R. Villumsen, I. Dalsgaard, L. Holten-Andersen, M.K. Raida, Potential role of specific antibodies as important vaccine induced protective mechanism against Aeromonas salmonicida in rainbow trout, PLoS One 7(10) (2012).

[35] R.N. Grøntvedt, S. Espelid, Vaccination and immune responses against atypical Aeromonas salmonicida in spotted wolffish (Anarhichas minor Olafsen) juveniles, Fish & Shellfish Immunology 16(3) (2004) 271–285.

[36] P. Vanden Bergh, M. Heller, S. Braga-Lagache, J. Frey, The Aeromonas salmonicida subsp. salmonicida exoproteome: global analysis, moonlighting proteins and putative antigens for vaccination against furunculosis, Proteome Science 11(1) (2013) 44.

[37] R.C. Cipriano, J. Bertolini, Selection for virulence in the fish pathogen Aeromonas salmonicida, using coomassie brilliant blue agar, Journal of Wildlife Diseases 24(4) (1988) 672–678.

[38] R.A. Garduño, A.R. Moore, G. Olivier, A.L. Lizama, E. Garduño, W.W. Kay, Host cell invasion and intracellular residence by Aeromonas salmonicida: role of the S-layer, Can J Microbiol 46(7) (2000) 660–8.

[39] G. Olivier, Virulence of Aeromonas salmonicida: Lack of Relationship with Phenotypic Characteristics, Journal of Aquatic Animal Health 2(2) (1990) 119–127.

[40] E.E. Ishiguro, W.W. Kay, T. Ainsworth, J.B. Chamberlain, R.A. Austen, J.T. Buckley, T.J. Trust, Loss of virulence during culture of Aeromonas salmonicida at high temperature, Journal of Bacteriology 148(1) (1981) 333–340.

[41] A.E. Ellis, A.S. Burrows, K.J. Stapleton, Lack of relationship between virulence of Aeromonas salmonicida and the putative virulence factors: A-layer, extracellular proteases and extracellular haemolysins, Journal of Fish Diseases 11(4) (1988) 309–323.

[42] S. Gulla, S. Bayliss, B. Björnsdóttir, I. Dalsgaard, O. Haenen, E. Jansson, U. McCarthy, F. Scholz, M. Vercauteren, D. Verner-Jeffreys, Biogeography of the fish pathogen Aeromonas salmonicida inferred by vapA genotyping, FEMS microbiology letters 366(7) (2019) fnz074.

[43] D. Basti, D. Bouchard, A. Lichtenwalner, Safety of Florfenicol in the Adult Lobster (Homarus americanus), Journal of Zoo and Wildlife Medicine 42(1) (2011) 131–133, 3.

[44] D.A. Bouchard, Investigating Present-day Health Issues of the American Lobster (Homarus americanus), (2018).

[45] J. Treasurer, T. Birkbeck, Pseudomonas anguilliseptica associated with mortalities in lumpfish (Cyclopterus lumpus L.) reared in Scotland, Bulletin of the European Association of Fish Pathologists 38 (2018) 222–224.

[46] A. Adams, Progress, challenges and opportunities in fish vaccine development, Fish & shellfish immunology 90 (2019) 210–214.

[47] C.W.E. Embregts, M. Forlenza, Oral vaccination of fish: Lessons from humans and veterinary species, Developmental & Comparative Immunology 64 (2016) 118–137.

[48] S. Gudmundsdóttir, S. Lange, B. Magnadóttir, B.K. Gudmundsdóttir, Protection against atypical furunculosis in Atlantic halibut, Hippoglossus hippoglossus (L.); comparison of a commercial furunculosis vaccine and an autogenous vaccine, Journal of Fish Diseases 26(6) (2003) 331–338.

[49] R. Nordmo, A. Ramstad, Variables affecting the challenge pressure of Aeromonas salmonicida and Vibrio salmonicida in Atlantic salmon (Salmo salar L.), Aquaculture 171(1) (1999) 1–12.

