## Supplemental tables and figures for "A commercial autogenous injection vaccine protects ballan wrasse (*Labrus bergylta*, Ascanius) against *Aeromonas salmonicida vapA* type V"

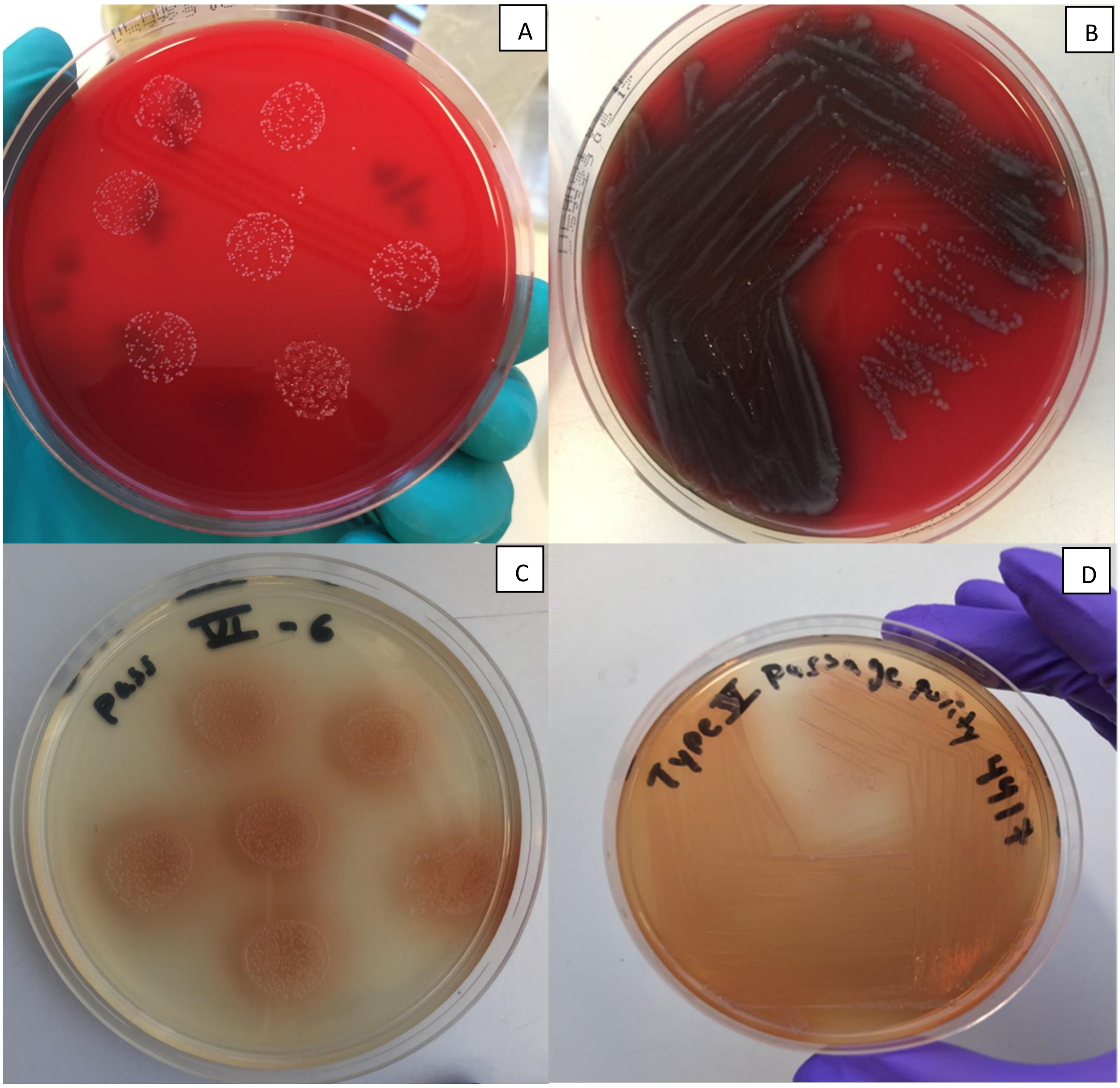


**Supplementary File 1.** Colony morphologyy of atypical *Aeromonas salmonicida* type VI**.** Coplonies were tiny and with a uniform size (A and C). Colonies produced a pigment which was more evident from day five post incubation (B, C and D) The pigment was grey in blood base agar (B) and brown in TSA.

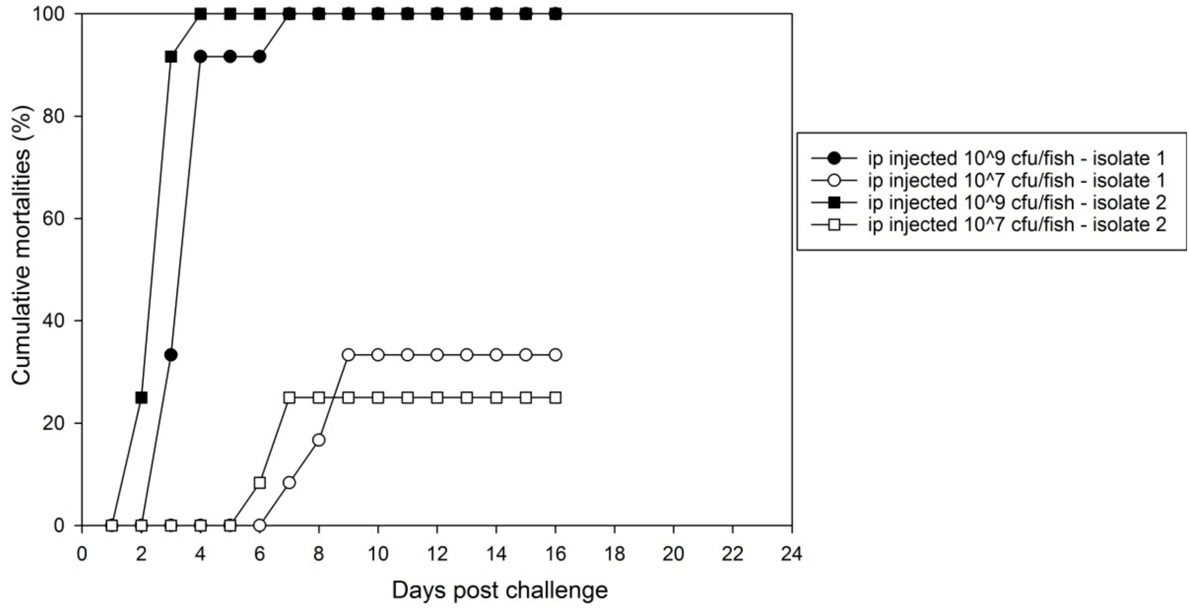


**Supplementary File 2.** Cumulative mortalities (%) of ballan wrasse (n= 12) challenged with different isolates of atypical *Aeromonas salmonicida* type VI at two infection doses. Isolate-1 TW164/15) and Isolate-2 TW184/16.

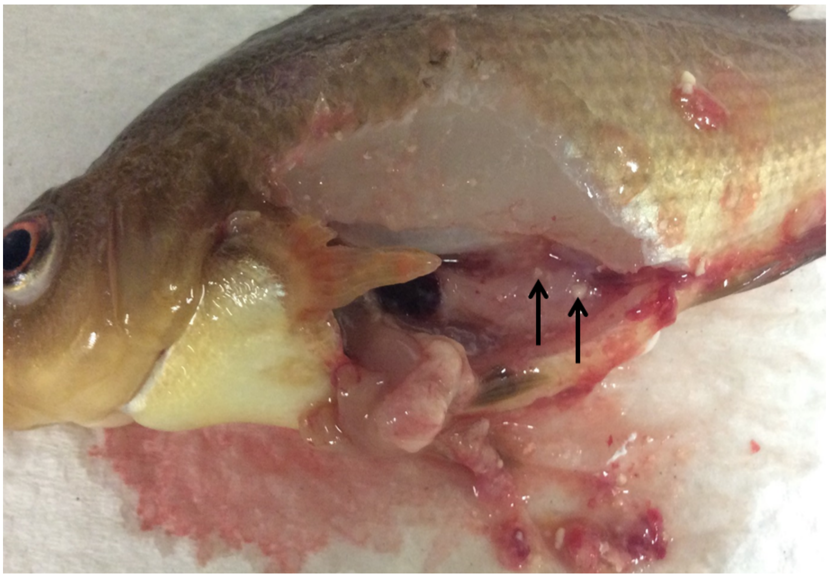

**Supplementary File 3.** Ballan wrasse experimentally infected with atypical *Aeromonas salmonicida* type VI (TW164/15). Extensive liquefaction of the organs and white nodules were present internally (black arrows).

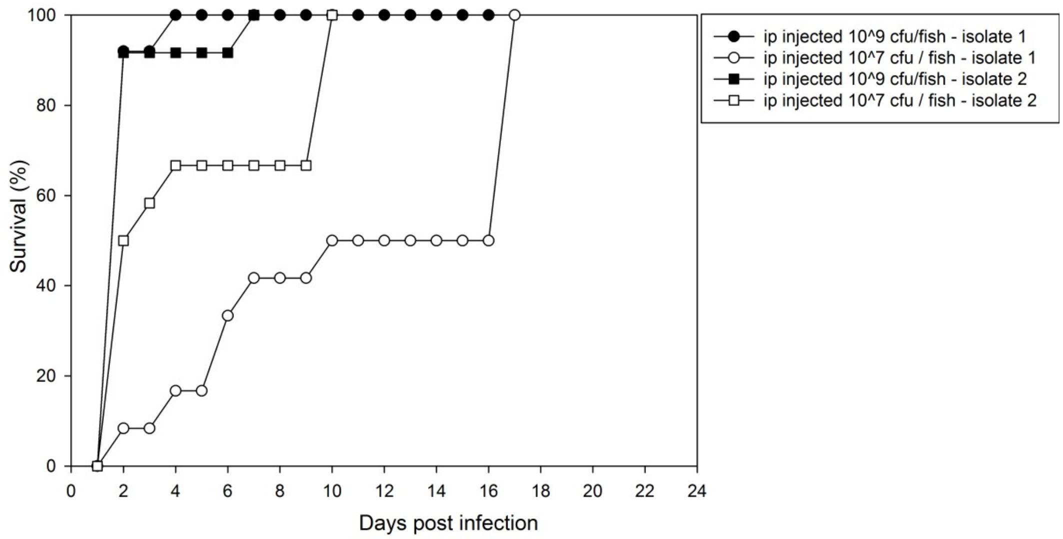


**Supplementary File 4.** Cumulative mortalities (%) of ballan wrasse (n= 12) challenged with different isolates of *Photobacterium indicum* at two injection doses. Isolate 1; TW138/16 and Isolate 2; 181/16.

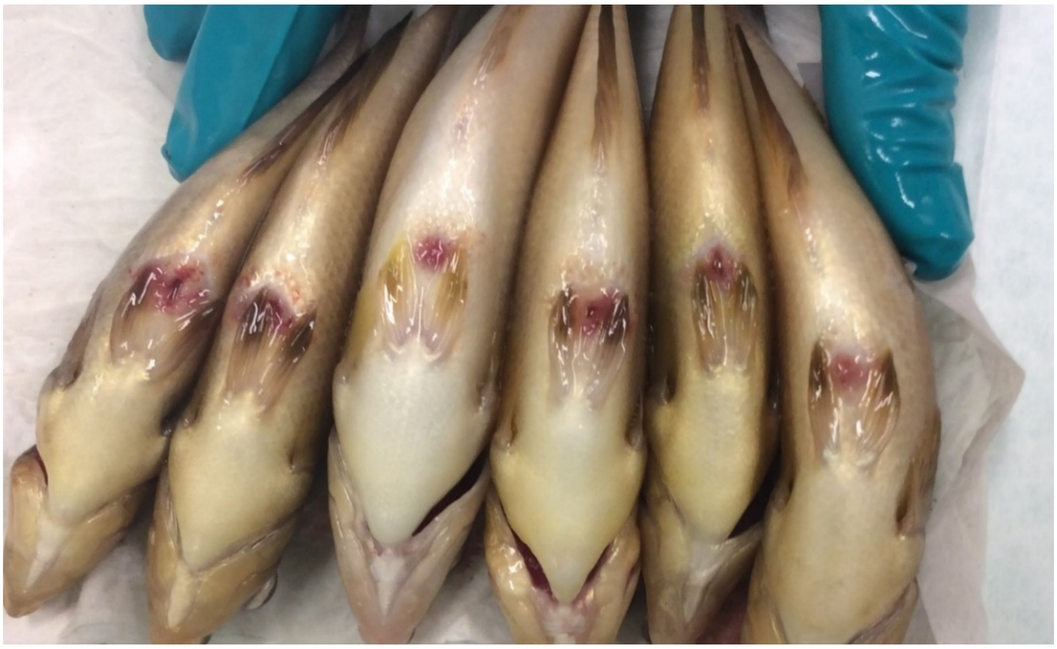

**Supplementary File 5.** Ballan wrasse survivors injected with *Photobacterium indicum* isolate (TW138/16) presenting ventral lesion around i.p. injection site at termination day (16 days post infection).

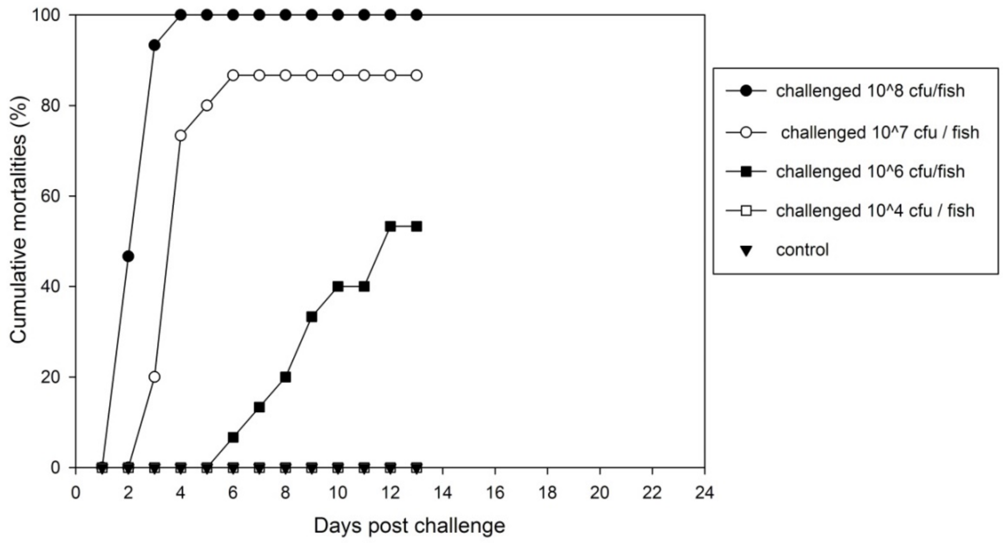


**Supplementary File 6.** Cumulative mortalities (%) of ballan wrasse challenged with atypical *Aeromonas salmonida* type V (TW4/14) using 4 doses; $10^8$, $10^7$, $10^6$ and $10^4$ cfu / fish (pre – test 1).

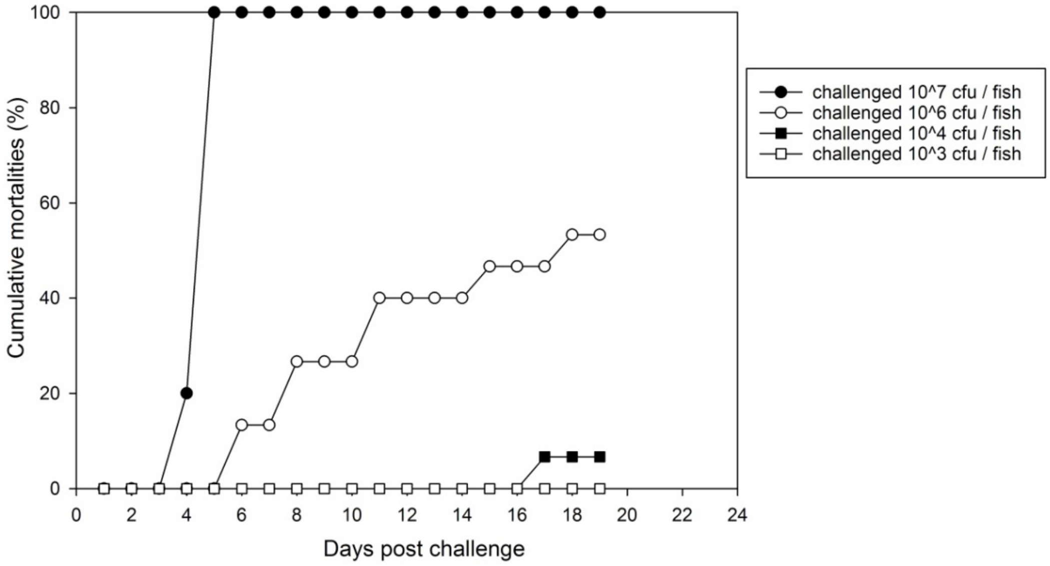


**Supplementary File 7.** Cumulative mortalities (%) of ballan wrasse challenged with atypical *Aeromonas salmonicida* type V (TW4/14) using four doses; $10^7$, $10^6$, $10^4$ and $10^3$ cfu / fish (pre – test 2).

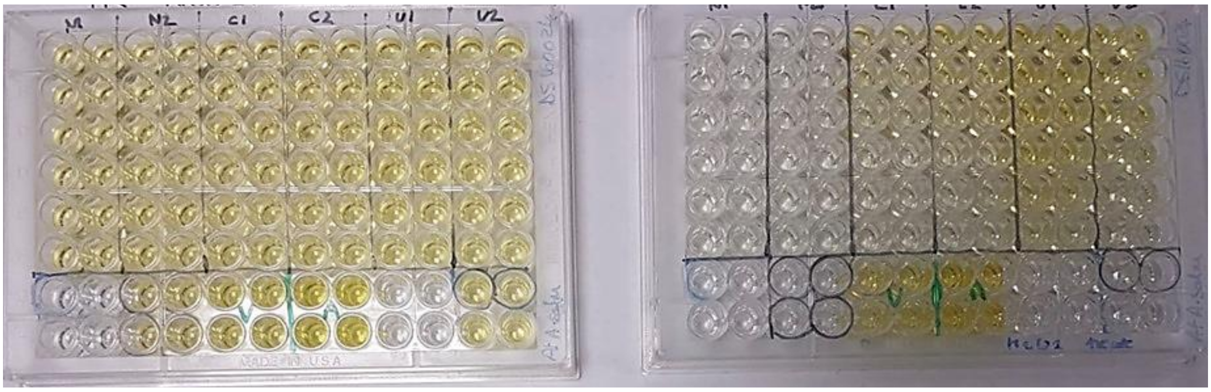

**Supplementary File 8.** ELISA optimisation for specific antibody (IgM) response to atypical *Aeromonas salmonicida* in ballan wrasse serum. ELISA 96-well plate with and without hydrogen peroxide blocking, left-hand side and right-hand side, respectively.

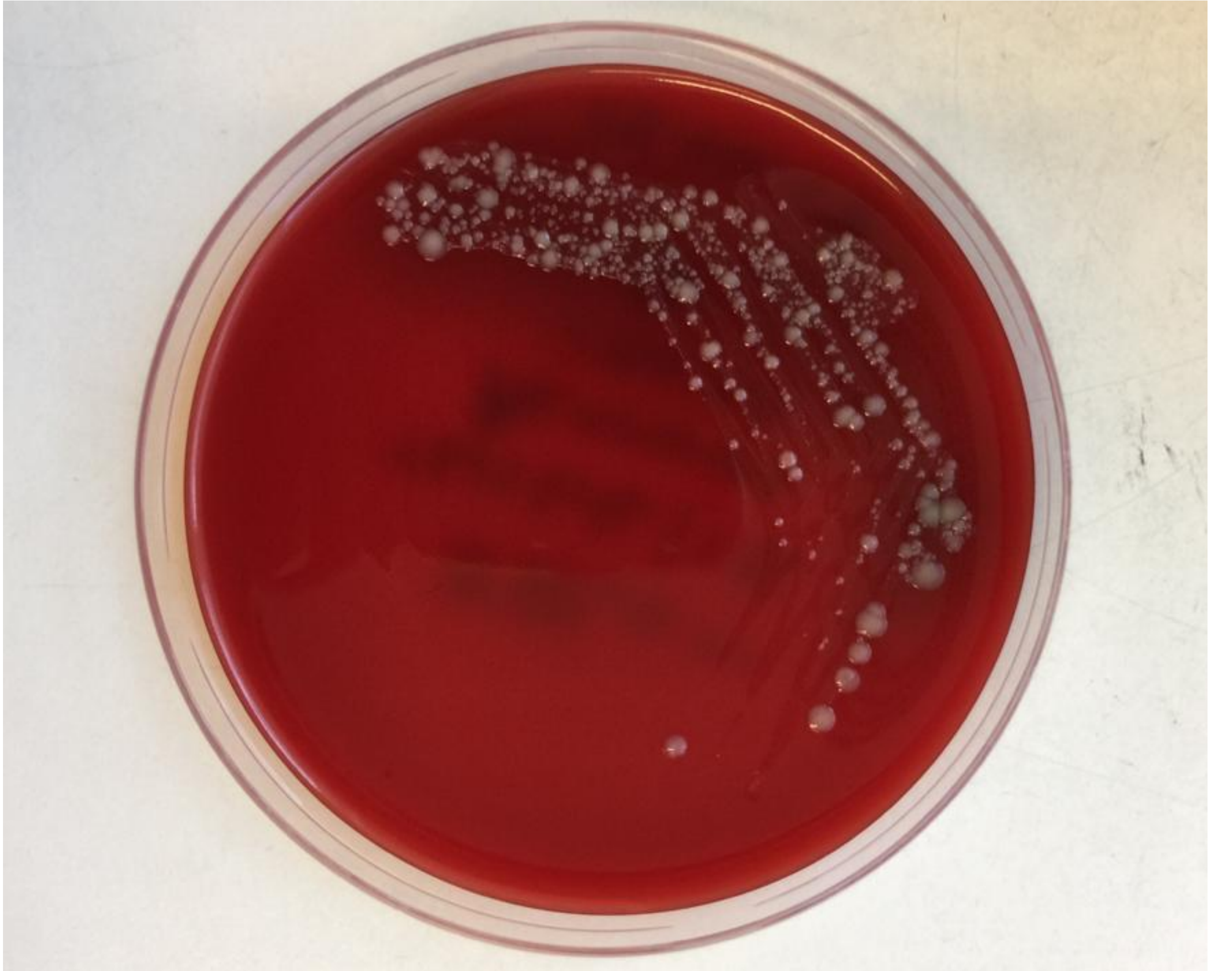

**Supplementary File 9.** Alternative colony morpholy of atypical *Aeromonas salmonicida* type VI. Colonies presented a mix of large and small size.
